# Force requirements of endocytic vesicle formation

**DOI:** 10.1101/2020.11.11.378273

**Authors:** Marc Abella, Lynnel Andruck, Gabriele Malengo, Michal Skruzny

## Abstract

Mechanical forces are integral to many cellular processes, including clathrin-mediated endocytosis, a principal membrane trafficking route into the cell. During endocytosis, forces provided by endocytic proteins and the polymerizing actin cytoskeleton reshape the plasma membrane into a vesicle. Assessing force requirements of endocytic membrane remodelling is essential for understanding endocytosis. Here, we determined forces applied during endocytosis using FRET-based tension sensors integrated into the major force-transmitting protein Sla2 in yeast. We measured force of approx. 10 pN transmitted over Sla2 molecule, hence a total force of 450-1300 pN required for endocytic vesicle formation. Importantly, decreasing cell turgor pressure and plasma membrane tension reduced force requirements of endocytosis. The measurements in hypotonic conditions and mutants lacking BAR-domain membrane scaffolds then showed the limits of the endocytic force-transmitting machinery. Our study provides force values and force profiles critical for understanding the mechanics of endocytosis and potentially other key cellular membrane-remodelling processes.

## Introduction

In clathrin-mediated endocytosis, the major trafficking pathway transporting nutrients, signals, pathogens and plasma membrane components into the cell, mechanical forces have to be applied to invaginate the plasma membrane and form endocytic vesicles (reviewed in Kaksonen and Roux, 2018; Lacy et al., 2018). These molecular forces are provided by the membrane-remodelling activities of endocytic proteins (e.g. ANTH, ENTH, or BAR domain proteins, mechanoenzyme dynamin), by formation of multiprotein scaffolds (e.g. clathrin coat), by protein crowding and by the polymerization of the actin cytoskeleton at the endocytic site (reviewed in Johannes et al., 2014; Stachowiak et al., 2013) The force supplied by actin polymerization is required when a high energy barrier of membrane deformation has to be overcome, like in cells with increased plasma membrane tension (Boulant et al., 2011; Kaplan et al., 2020) or high internal turgor pressure. In turgored cells, including yeasts, Arp2/3-mediated, branched actin polymerization is then an absolute prerequisite of membrane bending during endocytosis (Aghamohammadzadeh and Ayscough, 2009; Basu et al., 2014; Kukulski et al., 2012).

To transmit forces stored in actin polymerization and crosslinking, actin filaments have to be mechanically linked to the plasma membrane. This is accomplished by the conserved, force-transmitting protein linker made of endocytic adaptors Sla2-Ent1 in yeast and Hip1R-epsin 1-3 in human (Messa et al., 2014; Skruzny et al., 2012). These proteins bind cooperatively to the plasma membrane by their N-terminal ANTH and ENTH domains (Garcia-Alai et al., 2018; Lizarrondo et al., 2020; Skruzny et al., 2015) and redundantly to actin filaments through their C-terminal THATCH and ACB domains, respectively (Skruzny et al., 2012). The absence of Sla2/Hip1R-epsin linker completely blocks and severely impairs endocytosis in yeast and mouse cells, respectively (Messa et al., 2014; Skruzny et al., 2012).

The apparent full dependence of yeast endocytosis on force provided by the actin cytoskeleton and transmitted over the Sla2-Ent1 linker make this system uniquely suited for analyses of force requirements of endocytosis. We therefore aimed to measure forces applied on the Sla2-Ent1 linker during an endocytic event in yeast *Saccharomyces cerevisiae* using calibrated FRET-based tension sensor modules (TSMs). TSMs, which consist of two fluorophores undergoing efficient Förster resonance energy transfer (FRET) connected by a mechanosensitive peptide, reversibly extend at low pN force, reporting thus about applied forces by decrease in FRET (reviewed in Cost et al., 2019; Freikamp et al., 2017). Further, we sought to analyse roles of major environmental conditions and endocytic proteins in endocytic force transmission. We measured force of 450-1300 pN required for endocytic vesicle formation and show dependence of its magnitude on cellular turgor, plasma membrane tension and proper function of key membrane-remodelling endocytic factors.

## Results

### Sla2 force sensors report force required for endocytosis

To measure forces transmitted over the Sla2-Ent1 protein linker during endocytosis in yeast, we first constructed yeast strains expressing various Sla2 force sensors (Sla2 FS), Sla2 fusion proteins containing different TSMs. The TSMs, comprising of the efficient FRET pair mTurquoise2 and mNeonGreen (Mastop et al., 2017; Skruzny et al., 2019) connected by a peptide linker sensitive to distinct force ranges, F40 (1-6 pN), HP35 (6-8 pN) or HP35st (9-11 pN) (Austen et al., 2015; Grashoff et al., 2010), were inserted between the coiled-coil dimerization motif and the actin-binding THATCH domain of Sla2 (Fig. 1a). To distinguish force-dependent from force-independent FRET changes (Cost et al., 2019), we also constructed strains with all TSMs integrated after the THATCH domain, at the C-terminus of Sla2 (Sla2 no force controls, Sla2 NF; Supplementary Fig. 1). To channel force solely through Sla2 fusion proteins we also deleted the functionally redundant actin-binding domain of Ent1 in all strains (*ent1ΔACB* background; Skruzny et al., 2012). Resulting strains did not show any apparent defect in cell growth and endocytosis. Also, lifetimes and fluorescence intensities of Sla2 sensors at endocytic sites resemble the behaviour of wild-type, C-terminally-tagged Sla2 (Supplementary Fig. 1).

**Figure 1.**
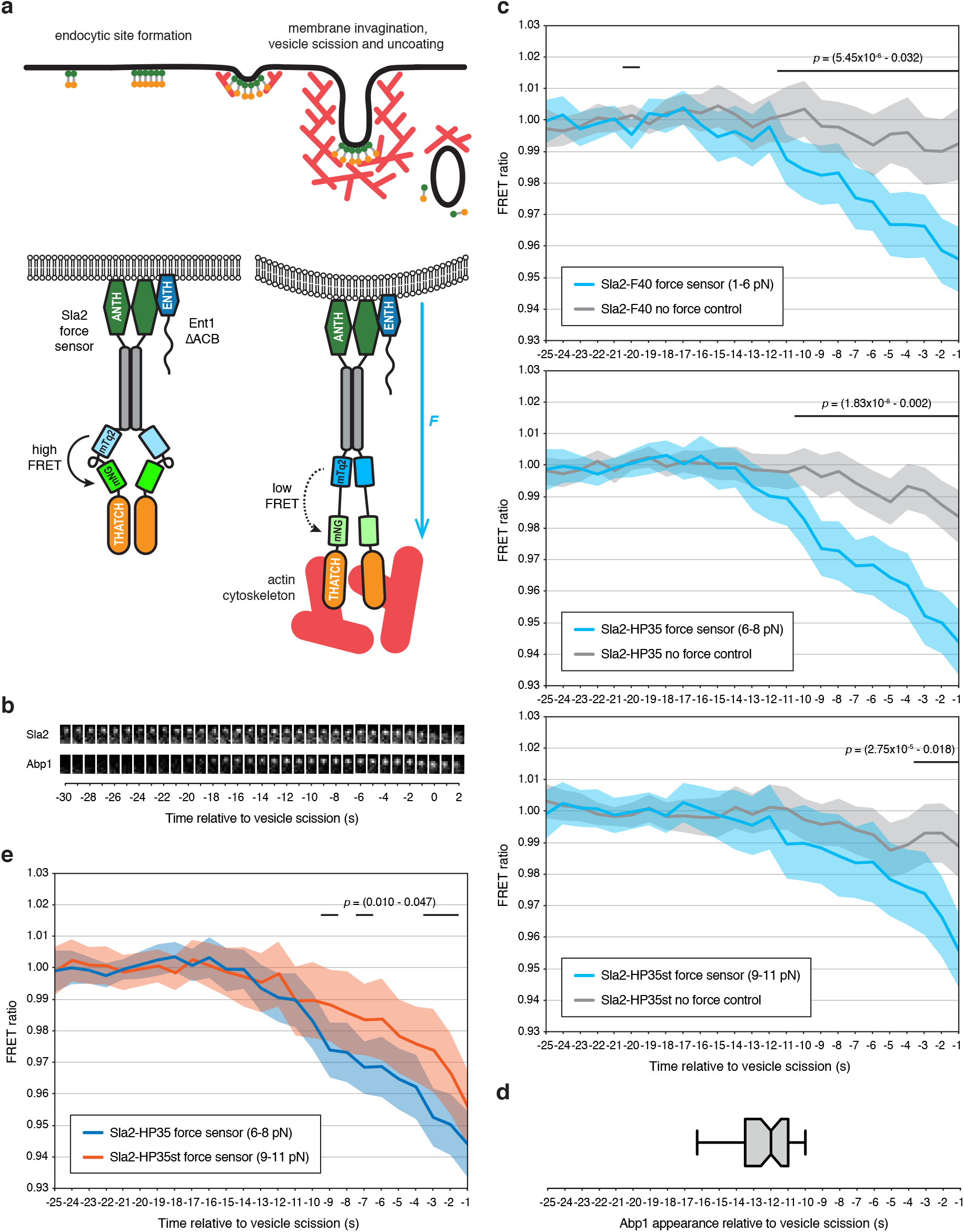
Force requirements of endocytic vesicle formation measured by Sla2 force sensors. **(a)** Timeline of endocytosis in yeast (top) and scheme of the Sla2 force sensor (bottom). (Top) For clarity, only Sla2 protein (green-orange rod) and actin filaments (red) are shown beneath the plasma membrane (black) though many other endocytic coat- and actin cytoskeleton-associated proteins participate in the process. (Bottom) FRET-based tension sensor module (TSM) consisting of mTurquoise2 (mTq2) and mNeonGreen (mNG) fluorophores connected by the mechanosensitive peptide (either F40, HP35, or HP35st) is inserted between the dimerization coiled-coil motif (grey bar) and the actin-binding THATCH domain of Sla2. Force applied on Sla2 sensor by polymerizing actin cytoskeleton causes peptide extension and a drop in FRET ratio between mNG and mTq2. All actin-supplied force is transmitted over the Sla2 sensor due to deletion of the redundant actin-binding domain of Ent1 (Ent1ΔACB). **(b)** Time series of fluorescence signals of Sla2-HP35 sensor (Sla2) and actin marker Abp1-mScarlet-I (Abp1) at the endocytic site. Tiles are oriented so that the cell exterior is up and the cell interior down. **(c)** FRET ratio profiles of Sla2 force sensors Sla2-F40 (1-6 pN; n=92), Sla2-HP35 (6-8 pN; n=108) and Sla2-HP35st (9-11 pN, n=93) (blue) and respective Sla2 no force controls (n=58, 82, 61; grey) acquired at individual endocytic sites before vesicle scission. Mean FRET ratios together with 95% confidence intervals are shown. Black lines indicate statistically significant differences between datasets with the range of p-values shown (Welch’s t-test). Force sensitivities of individual TSMs are indicated in parentheses. **(d)** Time of appearance of Abp1-mScarlet-I fluorescence signal at endocytic sites of Sla2-HP35 strain (n=45). Centre, right and left lines of the box plot indicate the median, and the 25th and 75th percentiles of the dataset, respectively. Whiskers show the 5th and 95th percentiles. **(e)** Comparison of FRET ratio profiles of Sla2-HP35 (blue; 6-8 pN) and Sla2-HP35st (red; 9-11 pN) sensors shown in **(c)**.

To follow the forces applied on the Sla2 force sensors, we analysed their FRET changes during individual endocytic events. Specifically, we simultaneously recorded their mTurquoise2 and mNeonGreen fluorescence signals at individual endocytic sites until the point of vesicle scission associated with a rapid move of the signal into the cytoplasm and its subsequent dissolution (Fig. 1b). Decrease of the ratio between mNeonGreen and mTurquoise2 fluorescence intensity (from here on FRET ratio) during endocytic events indicated force applied on all three Sla2 force sensors (Fig. 1c). Importantly, no changes in FRET ratio were observed for the respective Sla2 no force controls (Fig. 1c). Similarly, no change in FRET ratio was detected for strains with Sla2 force sensors lacking the actin-binding THATCH domain (Sla2ΔTHATCH NF controls) and channelling the force over the intact Ent1 protein (Supplementary Fig. 1). The absence of FRET changes in Sla2 NF and Sla2ΔTHATCH NF controls strongly suggests that the FRET ratio changes of Sla2 force sensors report on the force applied on Sla2 molecules and not on their conformational or intermolecular FRET changes potentially occurring during membrane invagination.

FRET ratio profiles of all three Sla2 force sensors showed an initial decrease in the FRET ratio approx. 13 s before vesicle scission (Fig. 1c). This timing coincided very well with the appearance of fluorescence signal of the actin marker Abp1-mScarlet-I at the endocytic sites (12.7 ± 2.6 s before vesicle scission; Fig. 1b,d), indicating that the force applied over the Sla2 sensors correlates with the onset of actin polymerization at the endocytic site (compare Fig. 1c and 1d). The similar starting point of the FRET ratio drop and its subsequent stepwise decrease, observed for all three sensors, furthermore suggests that Sla2 molecules are sequentially recruited to the growing actin cytoskeleton in course of endocytosis. When we compared FRET ratio profiles of different Sla2 force sensors, we found that while the Sla2-F40 and Sla2-HP35 sensors behaved similarly, a significantly smaller decrease in FRET ratio was observed for the Sla2-HP35st sensor in comparison to Sla2-HP35 (Fig. 1e). As the HP35st module is suited to detect higher forces (9-11 pN) than the structurally related HP35 module (6-8 pN; Austen et al., 2015), this indicates that not all Sla2-HP35st molecules are fully extended and the force applied over Sla2 therefore lies inside the dynamic range of HP35st, being approx. 10 pN. This value, in connection with the recently determined number of Sla2 molecules at the endocytic site (45-133 molecules; Picco et al., 2015; Sun et al., 2019), then results in approx. 450-1300 pN transmitted over Sla2 linker during a single endocytic event.

### Role of membrane-remodelling factors in endocytic force transmission

Having FRET-based endocytic force measurements established, we next analysed contributions of key membrane-remodelling factors to endocytic force transmission using the Sla2-HP35 sensor. First, we followed the Sla2-HP35 FRET ratio during so-called “retraction” endocytic events in cells deleted of the yeast amphiphysin homolog, BAR-domain protein Rvs167. Retraction events in a subset of endocytic sites of *rvs167Δ* cells are characterised by an initial slow movement of fluorescently-labelled endocytic marker towards the cytoplasm followed by its return back to the cell cortex, indicating that membrane invagination was aborted and the membrane retracted back to its initial flat conformation without actual vesicle scission (Fig. 2a; Kaksonen et al., 2005; Kishimoto et al., 2011). During retraction events, Sla2-HP35 FRET ratio first decreased to values typical for Sla2-HP35 sensor approx. 5 s before scission, and then it plateaued around these values, even though the actual fluorescence signal bounced back to the cell cortex (Fig. 2a). We conclude that similar forces are applied on the Sla2 linker in *rvs167Δ* cells during initial membrane bending and early invagination (and even on already retracted membrane), but the presence of Rvs167 protein is critical to facilitate their productive transmission during later invagination steps. This is in line with experimental and theoretical studies suggesting a key role of BAR-domain scaffolds in stabilization of long membrane profiles during endocytic invagination (Kishimoto et al., 2011; Kukulski et al., 2012; Walani et al., 2015). The values of maximal FRET ratio drop seen during retractions furthermore indicate that at least 50% of the total force transmitted over Sla2 is required for initial bending and shallow invagination of the plasma membrane.

**Figure 2.**
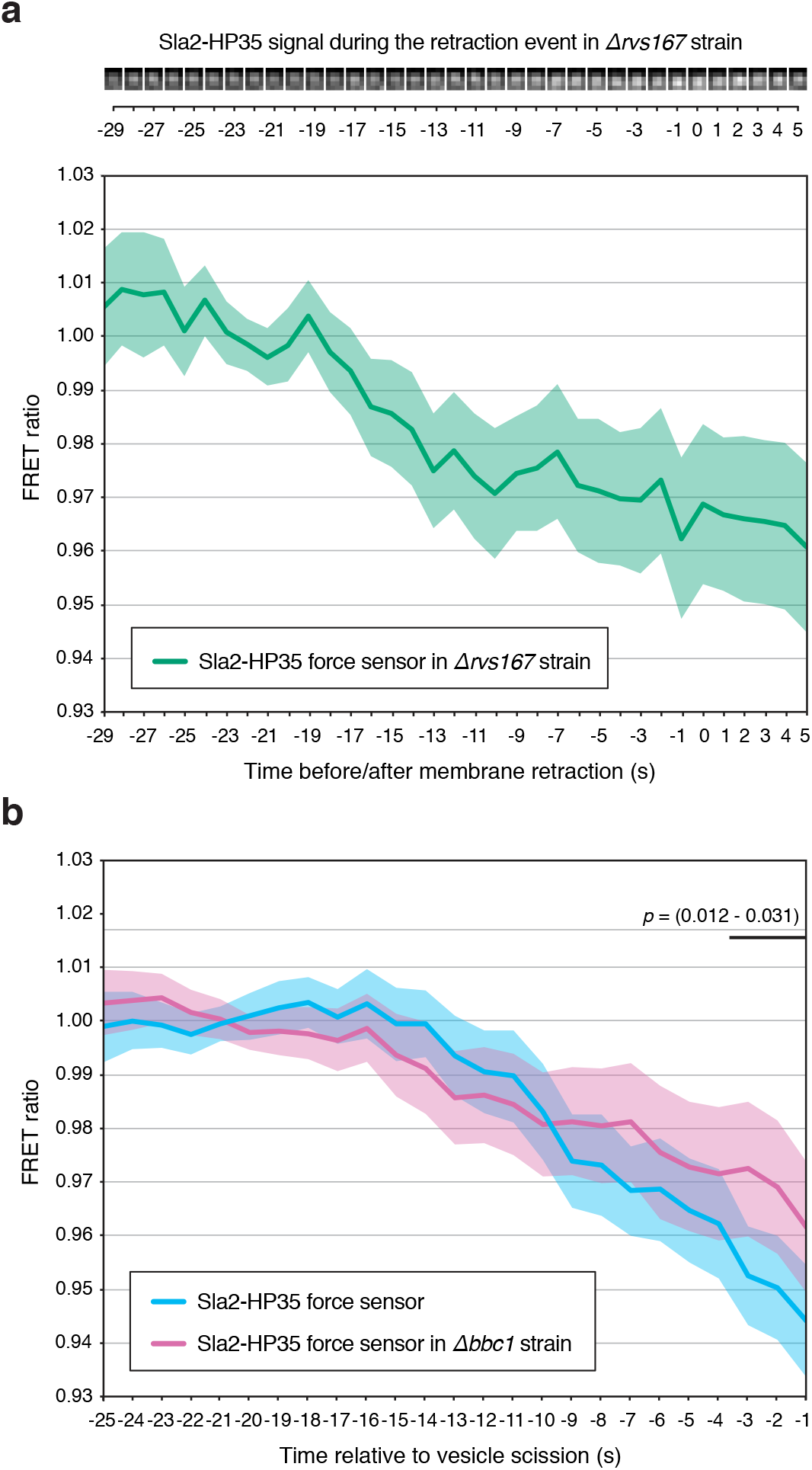
Role of membrane-remodelling factors in endocytic force transmission. **(a)** FRET ratio profile of Sla2-HP35 force sensor during endocytic retraction events in cells deleted of the amphiphysin homolog, BAR-domain protein Rvs167 (n=59). Time 0 is the time of the furthest move of Sla2-HP35 fluorescence signal in the cytoplasm before its retraction back to the cell cortex as illustrated by representative time series of Sla2-HP35 at the endocytic site at the top (oriented so that the cell exterior is up and the cell interior down). **(b)** FRET ratio profile of Sla2-HP35 sensor in cells deleted of Bbc1, negative regulator of actin polymerization at the endocytic site (purple; n=62). FRET ratio profile of Sla2-HP35 sensor in cells expressing wild-type Bbc1 protein (blue, as in Fig. 1c; n=108) is shown for comparison. Mean FRET ratio profiles together with 95% confidence intervals are shown. Black lines indicate statistically significant differences between datasets with the range of p-values shown (Welch’s t-test).

Next we analysed the role of organization of the actin cytoskeleton by following the Sla2-HP35 sensor in cells lacking protein Bbc1, a negative regulator of actin polymerization at the endocytic site (Kaksonen et al., 2005; Rodal et al., 2003). Although these cells have an enlarged endocytic actin cytoskeleton (Kaksonen et al., 2005; Picco et al., 2018), a smaller drop in the Sla2-HP35 FRET ratio was observed during the last 3 s before scission suggesting that less force is transmitted over Sla2 in the last phase of invagination (Fig. 2b). We propose that the extra force stored in the enlarged actin cytoskeleton of *bbc1Δ* cells directly remodel the long invaginating membrane e.g. by the expansion of the actin network suggested to happen a few seconds before vesicle scission (Picco et al., 2018).

### Decreased cell turgor pressure and plasma membrane tension reduce force requirements of endocytosis

With Sla2 force sensor system we then turned to test the effect of physical conditions on the force requirements of endocytosis. The huge internal turgor pressure of yeast cells (0.2-0.8 MPa; Goldenbogen et al., 2016; Schaber et al., 2010) represents the main mechanical barrier counteracting endocytic membrane invagination (Aghamohammadzadeh and Ayscough, 2009; Dmitrieff and Nédélec, 2015). We therefore tried to compensate for it by placing cells into hypertonic medium containing 0.25-0.5 M sorbitol, which should lower the osmotic difference between the cell cytoplasm and the surrounding environment. Remarkably, we detected a significantly smaller drop of Sla2-HP35 FRET ratio indicating that less force is transmitted over the sensor under the conditions reducing cellular turgor (Fig. 3a, Supplementary Fig. 2).

**Figure 3.**
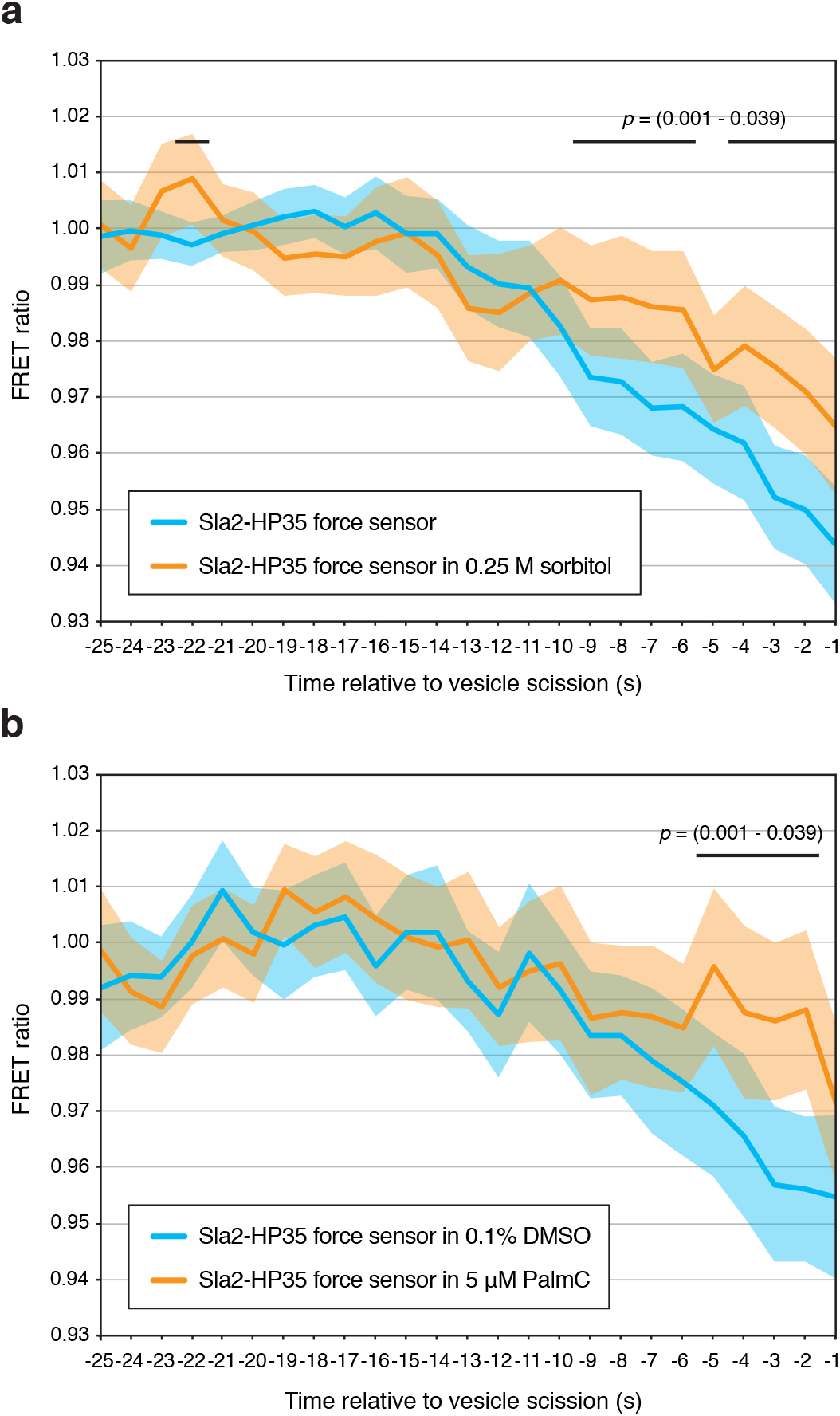
Decreased cell turgor pressure and plasma membrane tension reduce force requirements of endocytosis. **(a)** FRET ratio profile of Sla2-HP35 force sensor in cells incubated in medium with 0.25 M sorbitol for 5-15 min (orange; n=101). FRET ratio profile of Sla2-HP35 sensor in cells incubated without sorbitol (blue, as in Fig. 1c; n=108) is shown for comparison. **(b)** FRET ratio profiles of Sla2-HP35 sensor in cells incubated in medium containing 5 μM palmitoylcarnitine (PalmC) in 0.1% DMSO (orange; n=99) or 0.1% DMSO only (blue; n=87) for 35-45 min. Mean FRET ratio profiles together with 95% confidence intervals are shown. Black lines indicate statistically significant differences between datasets with the range of p-values shown (Welch’s t-test).

To corroborate these results by other means, we employed a recently described approach to reduce yeast plasma membrane tension without modifying the osmotic gradient across the plasma membrane (Riggi et al., 2018). For this, we incubated cells with soluble lipid palmitoylcarnitine (PalmC), which is incorporated into the yeast plasma membrane and reduces its tension. Again, we observed a significantly smaller decrease in the Sla2-HP35 FRET ratio, indicating a lower amount of force transmitted over the force sensor (Fig. 3b). In summary, these results suggest that a decrease of cell turgor pressure or plasma membrane tension efficiently reduce the force required for endocytosis.

### The endocytic force transmission system becomes insufficient under conditions of increased cell turgor

Finally, we analysed the capacity of the endocytic force-transmitting machinery by following Sla2 force sensors in cells incubated under hypotonic conditions, which should intensify cell turgor opposing endocytosis. For this we made use of *fps1Δ* cells, which lack aquaglyceroporin and therefore cannot adapt to hypoosmotic conditions by glycerol efflux (Tamás et al., 1999). We cultivated *fps1Δ* cells containing Sla2-HP35 or Sla2-HP35st sensors in medium containing 1 M sorbitol and exposed them to osmotic shifts made by an exchange to media of lower osmolarity containing 0.5 M, 0.4 M and 0.25 M sorbitol. In 1 M sorbitol medium, we observed normal levels of endocytosis with only slightly extended lifetime of Sla2 FS signal at endocytic sites (84% of endocytic events completed during 4 min; Sla2-HP35 lifetime 63.3 ± 3.5 s vs. 51.5 ± 3.7 s in medium without sorbitol; compare Fig. 4a and Supplementary Fig. 1). Contrary to that, cells shifted to 0.5 M sorbitol medium showed a clear increase in the number of stalled endocytic events with the lifetime extending 4 min (32% of events completed during 4 min; Fig. 4a). When we analysed the Sla2-HP35st force sensor (likely not fully extended during endocytosis under normal conditions) in completed endocytic events of shifted cells, we did not observe any significant difference in its FRET ratio profile in comparison to non-shifted cells (Supplementary Fig. 3). Osmotic shift to 0.4 M sorbitol medium further exacerbated the number of stalled endocytic events (22% of events completed; Fig. 4a). The Sla2-HP35st FRET ratio of completed endocytic events in this condition showed slightly bigger, yet not significantly decreased drop in comparison to non-shifted cells (Fig. 4b). Finally, in cells shifted to 0.25 M sorbitol medium, almost all endocytic events were stalled for longer than 4 min precluding the analysis of the FRET ratio (less than 1% of events completed, Fig. 4a). In summary, while increasing the osmotic difference between the cell and the environment led to an increasing number of stalled endocytic events, we could not detect a significant increase of force transmitted over the Sla2-HP35st sensor during completed endocytic events. This strongly indicates that there is no remarkable adaptation of the force-generating and/or transmitting system to hypoosmotic conditions. We propose that either the actin cytoskeleton can provide only a finite amount of force for membrane invagination or the connection between actin and the Sla2-Ent1 linker cannot sustain the extra force needed under these conditions. The latter possibility is in line with the recently observed uncoupling of the Sla2-Ent1 linker and the actin cytoskeleton at conditions of high plasma membrane tension (Riggi et al., 2019). We hypothesise that under hypotonic conditions (probably very common in natural yeast habitats) yeasts maintain the vital endocytic process primarily by modulation of internal turgor and not by direct regulation of endocytic force-generating or transmitting machinery. This idea is in good agreement with the presence of robust and fast-responding signalling and homeostatic pathways used by yeast to adapt to environmental osmotic changes (CWI and HOG MAPK pathways, TORC2 pathway, etc.).

**Figure 4.**
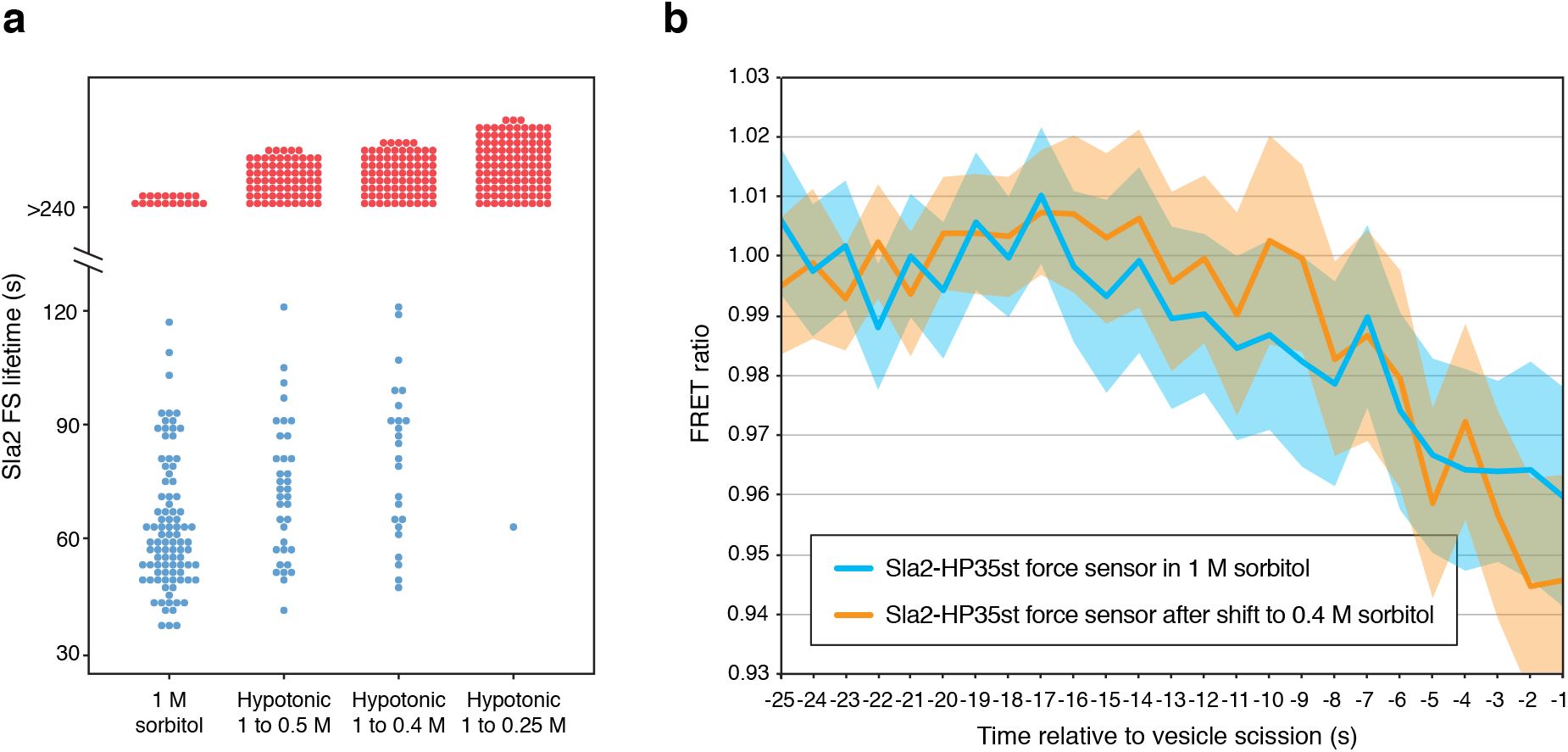
Endocytic force transmission system becomes insufficient under hypotonic conditions. **(a)** Lifetimes of fluorescence signal of Sla2-HP35 force sensor at endocytic sites of *fps1Δ* cells incubated in medium with 1 M sorbitol or shifted to hypotonic media with 0.5 M, 0.4 M and 0.25 M sorbitol for 30-60 min (blue dots). Endocytic sites arrested for the entire length of the 4 min acquisition are shown as red dots. (b) FRET ratio profiles of Sla2-HP35st sensor in *fps1Δ* cells incubated in 1 M sorbitol medium (blue; n=64) or shifted to medium with 0.4 M sorbitol for 30-60 min (orange; n=72). Mean FRET ratio profiles together with 95% confidence intervals are shown.

## Discussion

Many theoretical studies have attempted to estimate the forces required for endocytic membrane remodelling, proposing values over a range of two orders of magnitude (tens to thousands of pN; Akamatsu et al., 2020; reviewed in Carlsson, 2018; Lacy et al., 2018). Here, we have measured the force required for endocytic vesicle formation in yeast under normal and different genetically and environmentally perturbed conditions. Application of FRET-based Sla2 force sensors containing TSMs sensitive to different force ranges, together with the known character and number of force-transmitting molecules (Picco et al., 2015; Skruzny et al., 2012; Sun et al., 2019), allowed us to estimate the force provided by the actin cytoskeleton to be in the range of 450-1300 pN. We assume that these values are close to the upper limit of forces necessary for clathrin-mediated endocytosis in other organisms or cell types, as very high force has to be applied for endocytosis in yeasts to overcome their elevated turgor pressure.

However, recent studies indicate that substantial forces provided by the actin cytoskeleton are also necessary for endocytosis in the mammalian systems, e.g. in adherent cells or cells engulfing specific endocytic cargoes (Cureton et al., 2010; Hinze and Boucrot, 2018; Willy et al., 2017). The TSMs introduced into the homologous Hip1R-epsin 1-3 linker could be therefore used to study mechanics of endocytosis in these specialized cell types and tissues.

Knowledge of the force requirements of endocytic vesicle formation is not only essential for understanding how the endocytic machinery works in physiological and pathological conditions, but can also be useful for its biomedical and bioengineering applications (e.g. optimised drug delivery, selective molecular uptake). Finally, as force-driven remodelling of lipid membranes (e.g. by protein scaffolds, actin cytoskeleton, molecular motors) is central to many other cellular processes (Anitei and Hoflack, 2012; Jarsch et al., 2016), we anticipate that our approach can be pivotal for mechanobiological studies of other fundamental membrane-reshaping phenomena inside the cell.

## Materials and Methods

### Construction of FRET-based Sla2 force sensors

Plasmid pRS416-Sla2-HA (Skruzny et al., 2015) was used for assembling TSM sequences into respective positions of *SLA2* gene. First, *Sgr*AI and *Xba*I sites were inserted into *SLA2* gene after the codon for amino acid 702 (Sla2 FS and Sla2ΔTHATCH NF constructs) or before C-terminal HA-tag sequence (Sla2 NF constructs) by overlapping PCR (skipping the sequence coding for THATCH domain in case of Sla2ΔTHATCH NF construct). Similarly, overlapping PCR was used to construct TSM fragment containing mTurquoise2 and mNeonGreen sequences connected by sequence coding for F40 peptide and 5’-*Xma*I and 3’-*Xba*I overhangs. Both backbones and TSM insert were then cut by *Sgr*AI-*Xba*I and *Xma*I-*Xba*I, respectively, and ligated to give plasmids pRS416-Sla2(aa 1-702)-mTurquoise2-F40-mNeonGreen-Sla2(aa 703-968) alias Sla2-F40 FS; pRS416-Sla2(aa 1-702)-mTurquoise2-F40-mNeonGreen alias Sla2ΔTHATCH-F40 NF; and pRS416-Sla2-mTurquoise2-F40-mNeonGreen alias Sla2-F40 NF. Sequences coding HP35 and HP35st linkers with *Spe*I and *Age*I overhangs were synthetised commercially (IDT) and exchanged with F40 sequence (containing *Spe*I and *Age*I sites generated by codon optimization) in above plasmids by *Spe*I-*Age*I digest and ligation. Resulting plasmids were amplified by PCR with primers covering *SLA2* fragment with TSM module, terminator sequence and *URA3* locus of the plasmid. The reverse primer also contained 63 bp overhang homologous to 3’ UTR of *SLA2* gene. These PCR fragments were integrated to genomic *SLA2* locus using standard lithium acetate transformation. Correct integration was confirmed by sequencing. In all Sla2 force sensors and no force controls the Sla2 protein thus contains following insertion: Thr-Gly-mTurquoise2-Thr-Ser-TSM sequence-Thr-Gly-mNeonGreen-Ser-Arg at positions indicated above.

### Yeast strains and media

Standard yeast media and protocols were used to manipulate yeast strains listed in Supplementary Table 1. The C-terminal tagging and deletion of yeast proteins was made by homologous recombination of respective genes with PCR cassettes previously described (Janke et al., 2004) or constructed in the lab. For microscopy, strains were grown to a log phase in a low fluorescence SD-Trp, -Ura medium (prepared from LoFlo YNB; Formedium) containing 1 M sorbitol where indicated. Cells were attached to Concanavalin A-coated (0.1 μg/ml; Sigma-Aldrich) 8-well glass slides (ibidi) and observed at 20 °C. Changes in the medium composition were performed by replacing initial medium with medium containing sorbitol, palmitoylcarnitine (PalmC; Sigma-Aldrich) in DMSO or DMSO only as indicated in the text.

### Microscopy

Ratiometric FRET imaging was performed using a wide-field Eclipse Ti-E fluorescence microscope (Nikon) equipped with X-Cite Exacte LED light source, Perfect Focus System (PFS) and NIS-Elements AR software (4.40; Nikon). Typically, 75 images with 1 s frame rate were acquired with Nikon 100x Plan Apo λ, NA 1.45 oil immersion objective and iXon 897-X3 EM-CCD camera (Andor) with EM gain set up to 250. The excitation filter (ET436/20x; Chroma) was used to excite mTurquoise2 FRET donor, and both donor and mNeonGreen FRET acceptor fluorescence were simultaneously detected by the camera using a fluorescence splitting device (Optosplit II; CAIRN) containing emission filters (ET480/40m; Chroma) and (FF01-542/27; Semrock) and FF520-Di02 beam splitter (Semrock).

To compare behaviour of Sla2 force sensors, Sla2-mNeonGreen and Abp1-mScarlet-I at endocytic sites, 4 min movies were acquired with 1 s frame rate by filters set FF01-504/12 (excitation), FF520-Di02 (dicroic) and FF01-542/27 (emission) for mNeonGreen, and with 0.5 s frame rate by FF01-562/40 (excitation), FF605-Di02 (dichroic) and FF01-647/57 (emission; all Semrock) for mScarlet-I fluorescence. Images were analysed with FIJI software (Schindelin et al., 2012). The mTurquoise2 and mNeonGreen images were first aligned, subtracted of general background and corrected for photobleaching, and individual endocytic events were then manually tracked using TrackMate plugin (Tivenez et al., 2017). Time alignment of individual endocytic events was performed taking vesicle scission as reference point (time 0 s). Only tracks with minimally 20 consecutive time points before the scission were analysed, including up to 35 track points into the analysis (see Supplementary Tables 2-13). FRET ratio of individual tracks was normalised by the average FRET ratio value of track points −25 to −17 s, where mTurquoise2 and mNeonGreen FRET ratio was largely unchanged. In total 58-121 endocytic tracks were analysed to obtain the average FRET ratio profile. All FRET ratio data are summarised in Supplementary Tables 2-13.

### Statistics and reproducibility

The samples sizes were based on previous quantitative fluorescence microscopy studies of yeast endocytosis (Galletta et al., 2008; Picco et al., 2015; Sun et al., 2019) and protocols for TSM microscopy (Cost et al., 2019). Fluorescence data from 58-121 endocytic patches (of approx. 55-115 cells) acquired during 3-5 independent imaging sessions were analysed for individual sensors and conditions as indicated in figure legends. Two-tailed Welch’s t-test for unpaired datasets of uneven variances was used to compare individual time points of Sla2 force sensors with the same time points of respective Sla2 control datasets, or with time points of Sla2 force sensors acquired under indicated conditions.

## Acknowledgements

We thank M.-L. Hocke for her help with TrackMate analysis; M. Kaksonen (University of Geneva) for initial discussions about the project, C. Grashoff (University of Münster) for his kind advice on tension sensor modules; and J. Freitag (University of Marburg) and S.M. Murray (MPI Marburg) for critical reading of the manuscript. This work was funded by Deutsche Forschunsgemeinschaft (DFG) Research Grant SK 305/1-1.

## Author contributions

M.S. conceived the work together with M.A. M.A. developed the methods together with M.S. and G.M. M.A. performed and analysed all experiments together with contribution of L.A. and M.S. M.S. wrote the manuscript with the input from M.A. and G.M., supervised the project and secured funding.

## References

Aghamohammadzadeh, S., and Ayscough, K.R. (2009). Differential requirements for actin during yeast and mammalian endocytosis. Nat. Cell Biol. 11, 1039–1042.

Akamatsu, M., Vasan, R., Serwas, D., Ferrin, M., Rangamani, P., and Drubin, D.G. (2020). Principles of self-organization and load adaptation by the actin cytoskeleton during clathrin-mediated endocytosis. Elife 9, 1–40.

Anitei, M., and Hoflack, B. (2012). Bridging membrane and cytoskeleton dynamics in the secretory and endocytic pathways. Nat. Cell Biol. 14, 11–19.

Austen, K., Ringer, P., Mehlich, A., Chrostek-Grashoff, A., Kluger, C., Klingner, C., Sabass, B., Zent, R., Rief, M., and Grashoff, C. (2015). Extracellular rigidity sensing by talin isoform-specific mechanical linkages. Nat. Cell Biol. 17, 1597–1606.

Basu, R., Munteanu, E.L., and Chang, F. (2014). Role of turgor pressure in endocytosis in fission yeast. Mol. Biol. Cell 25, 679–687.

Boulant, S., Kural, C., Zeeh, J.C., Ubelmann, F., and Kirchhausen, T. (2011). Actin dynamics counteract membrane tension during clathrin-mediated endocytosis. Nat. Cell Biol. 13, 1124–1132.

Carlsson, A.E. (2018). Membrane bending by actin polymerization. Curr. Opin. Cell Biol. 50, 1–7.

Cost, A.L., Khalaji, S., and Grashoff, C. (2019). Genetically Encoded FRET-Based Tension Sensors. Curr. Protoc. Cell Biol. 83, 1–29.

Cureton, D.K., Massol, R.H., Whelan, S.P.J., and Kirchhausen, T. (2010). The length of vesicular stomatitis virus particles dictates a need for actin assembly during clathrin-dependent endocytosis. PLoS Pathog. 6.

Dmitrieff, S., and Nédélec, F. (2015). Membrane Mechanics of Endocytosis in Cells with Turgor. PLoS Comput. Biol. 11, 1–15.

Freikamp, A., Mehlich, A., Klingner, C., and Grashoff, C. (2017). Investigating piconewton forces in cells by FRET-based molecular force microscopy. J. Struct. Biol. 197, 37–42.

Galletta, B.J., Chuang, D.Y., and Cooper, J.A. (2008). Distinct roles for Arp2/3 regulators in actin assembly and endocytosis. PLoS Biol. 6, 0072–0085.

Garcia-Alai, M.M., Heidemann, J., Skruzny, M., Gieras, A., Mertens, H.D.T., Svergun, D.I., Kaksonen, M., Uetrecht, C., and Meijers, R. (2018). Epsin and Sla2 form assemblies through phospholipid interfaces. Nat. Commun. 9.

Goldenbogen, B., Giese, W., Hemmen, M., Uhlendorf, J., Herrmann, A., and Klipp, E. (2016). Dynamics of cell wall elasticity pattern shapes the cell during yeast mating morphogenesis. Open Biol. 6.

Grashoff, C., Hoffman, B.D., Brenner, M.D., Zhou, R., Parsons, M., Yang, M.T., McLean, M.A., Sligar, S.G., Chen, C.S., Ha, T., et al. (2010). Measuring mechanical tension across vinculin reveals regulation of focal adhesion dynamics. Nature 466, 263–266.

Hinze, C., and Boucrot, E. (2018). Endocytosis in proliferating, quiescent and terminally differentiated cells. J. Cell Sci. 131, 1–10.

Janke, C., Magiera, M.M., Rathfelder, N., Taxis, C., Reber, S., Maekawa, H., Moreno-Borchart, A., Doenges, G., Schwob, E., Schiebel, E., et al. (2004). A versatile toolbox for PCR-based tagging of yeast genes: New fluorescent proteins, more markers and promoter substitution cassettes. Yeast 21, 947–962.

Jarsch, I.K., Daste, F., and Gallop, J.L. (2016). Membrane curvature in cell biology: An integration of molecular mechanisms. J. Cell Biol. 214, 375–387.

Johannes, L., Wunder, C., and Bassereau, P. (2014). Bending “on the rocks”-A cocktail of biophysical modules to build endocytic pathways. Cold Spring Harb. Perspect. Biol. 6, 1–17.

Kaksonen, M., and Roux, A. (2018). Mechanisms of clathrin-mediated endocytosis. Nat. Rev. Mol. Cell Biol. 19, 313–326.

Kaksonen, M., Toret, C.P., and Drubin, D.G. (2005). A modular design for the clathrin- and actin-mediated endocytosis machinery. Cell 123, 305–320.

Kaplan, C., Kenny, S.J., Chen, S., Sitarska, E., Xu, K., Drubin, D.G., Biology, C., Biology, C., Unit, B., Molecular, E., et al. (2020). Short running title : Adaptive actin networks ensure robust endocytosis Full running title : Adaptive actin organization counteracts elevated membrane tension to ensure robust endocytosis Highlights : Summary (~ 40-word): 664.

Kishimoto, T., Sun, Y., Buser, C., Liu, J., Michelot, A., and Drubin, D.G. (2011). Determinants of endocytic membrane geometry, stability, and scission. Proc. Natl. Acad. Sci. U. S. A. 108.

Kukulski, W., Schorb, M., Kaksonen, M., and Briggs, J.A.G. (2012). Plasma membrane reshaping during endocytosis is revealed by time-resolved electron tomography. Cell 150, 508–520.

Lacy, M.M., Ma, R., Ravindra, N.G., and Berro, J. (2018). Molecular mechanisms of force production in clathrin-mediated endocytosis. FEBS Lett. 592, 3586–3605.

Lizarrondo, J., Klebl, D.P., Niebling, S., Abella, M., Schroer, M.A., Mertens, H.D.T., Veith, K., Svergun, D.I., Skruzny, M., Sobott, F., et al. Structure of the endocytic adaptor complex reveals the basis for efficient membrane anchoring during clathrin-mediated endocytosis.

Mastop, M., Bindels, D.S., Shaner, N.C., Postma, M., Gadella, T.W.J., and Goedhart, J. (2017). Characterization of a spectrally diverse set of fluorescent proteins as FRET acceptors for mTurquoise2. Sci. Rep. 7, 1–18.

Messa, M., Fernández-Busnadiego, R., Sun, E.W., Chen, H., Czapla, H., Wrasman, K., Wu, Y., Ko, G., Ross, T., Wendland, B., et al. (2014). Epsin deficiency impairs endocytosis by stalling the actin-dependent invagination of endocytic clathrin-coated pits. Elife 3, 1–25.

Picco, A., Mund, M., Ries, J., Nédélec, F., and Kaksonen, M. (2015). Visualizing the functional architecture of the endocytic machinery. Elife 2015, 1–29.

Picco, A., Kukulski, W., Manenschijn, H.E., Specht, T., Briggs, J.A.G., and Kaksonen, M. (2018). The contributions of the actin machinery to endocytic membrane bending and vesicle formation. Mol. Biol. Cell 29, 1346–1358.

Riggi, M., Niewola-Staszkowska, K., Chiaruttini, N., Colom, A., Kusmider, B., Mercier, V., Soleimanpour, S., Stahl, M., Matile, S., Roux, A., et al. (2018). Decrease in plasma membrane tension triggers PtdIns(4,5)P 2 phase separation to inactivate TORC2. Nat. Cell Biol. 20, 1043–1051.

Riggi, M., Bourgoint, C., Macchione, M., Matile, S., Loewith, R., and Roux, A. (2019). TORC2 controls endocytosis through plasma membrane tension. J. Cell Biol. 218, 2265–2276.

Rodal, A.A., Manning, A.L., Goode, B.L., and Drubin, D.G. (2003). Negative Regulation of Yeast WASp by Two SH3 Domain-Containing Proteins. Curr. Biol. 13, 1000–1008.

Schaber, J., Adrover, M.À., Eriksson, E., Pelet, S., Petelenz-Kurdziel, E., Klein, D., Posas, F., Goksör, M., Peter, M., Hohmann, S., et al. (2010). Biophysical properties of Saccharomyces cerevisiae and their relationship with HOG pathway activation. Eur. Biophys. J. 39, 1547–1556.

Schindelin, J., Arganda-Carreras, I., Frise, E., Kaynig, V., Longair, M., Pietzsch, T., Preibisch, S., Rueden, C., Saalfeld, S., Schmid, B., et al. (2012). Fiji: An opensource platform for biological-image analysis. Nat. Methods 9, 676–682.

Skruzny, M., Brach, T., Ciuffa, R., Rybina, S., Wachsmuth, M., and Kaksonen, M. (2012). Molecular basis for coupling the plasma membrane to the actin cytoskeleton during clathrin-mediated endocytosis. Proc. Natl. Acad. Sci. U. S. A. 109, E2533–42.

Skruzny, M., Desfosses, A., Prinz, S., Dodonova, S.O., Gieras, A., Uetrecht, C., Jakobi, A.J., Abella, M., Hagen, W.J.H., Schulz, J., et al. (2015). An Organized Co-assembly of Clathrin Adaptors Is Essential for Endocytosis. Dev. Cell 33.

Skruzny, M., Pohl, E., and Abella, M. (2019). FRET microscopy in yeast. Biosensors.

Stachowiak, J.C., Brodsky, F.M., and Miller, E.A. (2013). A cost-benefit analysis of the physical mechanisms of membrane curvature. Nat. Cell Biol. 15, 1019–1027.

Sun, Y., Schöneberg, J., Chen, X., Jiang, T., Kaplan, C., Xu, K., Pollard, T.D., and Drubin, D.G. (2019). Direct comparison of clathrin-mediated endocytosis in budding and fission yeast reveals conserved and evolvable features. Elife 8.

Tamás, M.J., Luyten, K., Sutherland, F.C.W., Hernandez, A., Albertyn, J., Valadi, H., Li, H., Prior, B.A., Kilian, S.G., Ramos, J., et al. (1999). Fps1p controls the accumulation and release of the compatible solute glycerol in yeast osmoregulation. Mol. Microbiol. 31, 1087–1104.

Tinevez, J.Y., Perry, N., and Schindelin, J. et al. (2017). TrackMate: An open and extensible platform for single-particle tracking. Methods 115, 80–90.

Walani, N., Torres, J., and Agrawal, A. (2015). Endocytic proteins drive vesicle growth via instability in high membrane tension environment. Proc. Natl. Acad. Sci. U. S. A. 112, E1423–E1432.

Willy, N.M., Ferguson, J.P., Huber, S.D., Heidotting, S.P., Aygün, E., Wurm, S.A., Johnston-Halperin, E., Poirier, M.G., and Kural, C. (2017). Membrane mechanics govern spatiotemporal heterogeneity of endocytic clathrin coat dynamics. Mol. Biol. Cell 28, 3480–3488.

