## Supplementary Information for "Force requirements of endocytic vesicle formation"

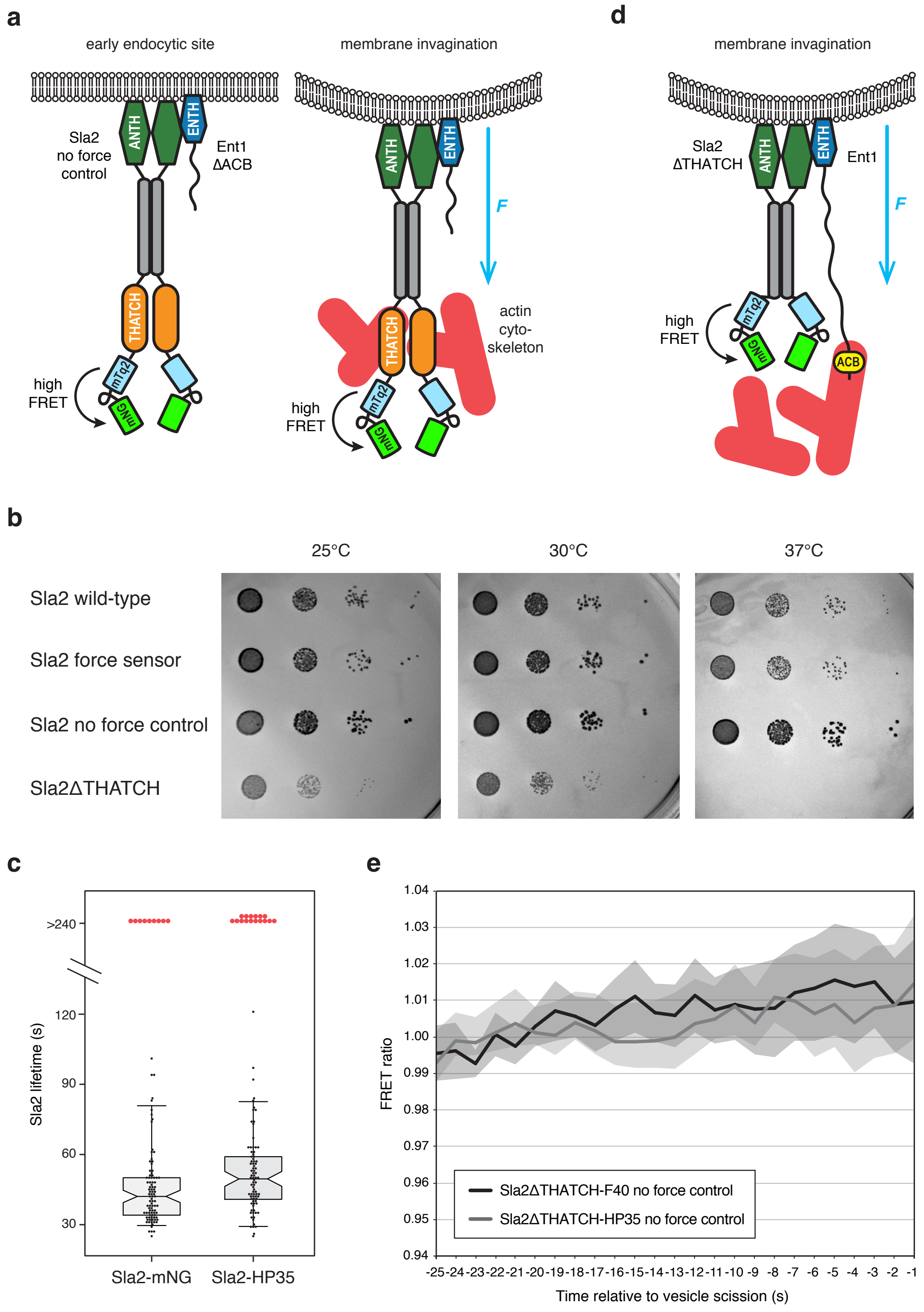


**Supplementary Figure 1** Functionality of the Sla2 FRET-based system to measure forces required for endocytosis. **(a)** Scheme of Sla2 no force sensor. FRET-based tension sensor module consisting of mTurquoise2 (mTq2) and mNeon-Green (mNG) fluorophores connected by the mechanosensitive peptide (either F40, HP35 or HP35st) is inserted after the actin-binding THATCH domain of Sla2 protein. Force applied on the Sla2 no force control by the actin cytoskeleton cannot therefore extend the mechanosensitive peptide and eventual FRET changes are not related to the transmission of force over Sla2. **(b)** Sla2 sensors complement function of Sla2 protein. Ten-fold serial dilutions of *sla2Δ, ent1ΔACB* strain expressing indicated proteins from *URA3 CEN* plasmid were incubated at indicated temperatures on SD-Ura plates for 1.5-2 days. Sla2ΔTHATCH construct unable to bind actin was used as negative control. **(c)** Lifetimes of fluorescence signals of Sla2-mNG (n=102) and Sla2-HP35 force sensor (n=100) proteins at endocytic sites before vesicle scission. Centre, bottom and top lines of box plots indicate the median, and the 25th and 75th percentiles of the datasets, respectively. Whiskers show the 5th and 95th percentiles. Signals persisting for the entire length of the 4 min acquisition are shown as red dots. Imaging settings were identical for both constructs. **(d)** Scheme of Sla2ΔTHATCH no force sensor. TSM is inserted after the dimerization coiled-coil motif (grey bar) of Sla2 protein lacking of the actin-binding THATCH domain. All actin-supplied force is transmitted over the actin-binding ACB domain of Ent1. Eventual FRET changes of Sla2ΔTHATCH sensor are thus not related to the force transmission over Sla2. **(e)** FRET ratio profiles of Sla2ΔTHATCH-F40 (dark grey; n=26) and Sla2ΔTHATCH-HP35 (light grey; n=31) no force controls acquired at endocytic sites before vesicle scission. Mean FRET ratio profiles together with 95% confidence intervals are shown.

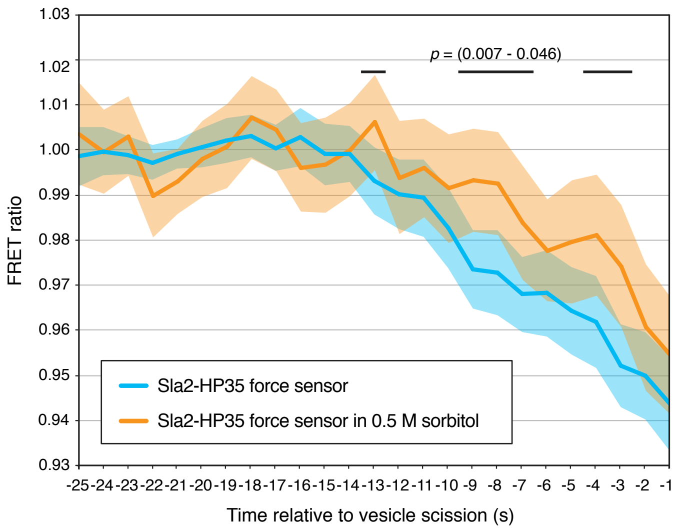


**Supplementary Figure 2** Decreased cell turgor pressure reduces force requirements of endocytosis. FRET ratio profile of Sla2-HP35 force sensor in cells incubated in medium with 0.5 M sorbitol for 5-15 min (orange; n=80). FRET ratio profile of Sla2-HP35 sensor in cells incubated without sorbitol (blue, as in Fig. 1c; n=108) is shown for comparison. Mean FRET ratio profiles together with 95% confidence intervals are shown. Black lines indicate statistically significant differences between datasets with the range of p-values shown (Welch's t-test).

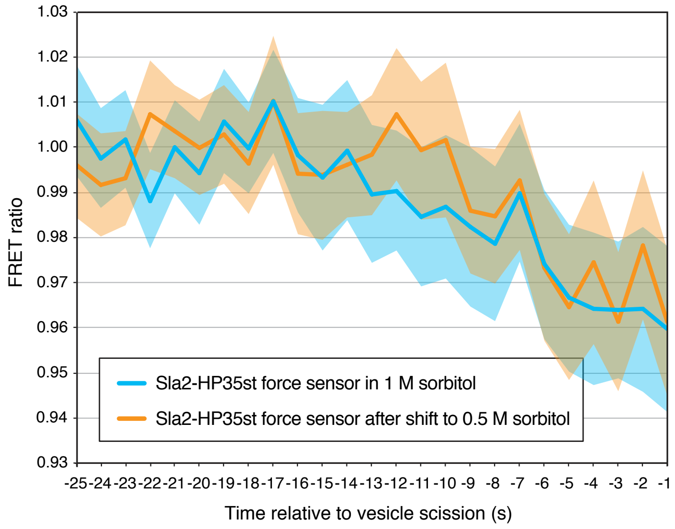


**Supplementary Figure 3** Endocytic force transmission system becomes insufficient under hypotonic conditions. FRET ratio profiles of Sla2-HP35st sensor in *fps1Δ* cells incubated in 1 M sorbitol medium (blue; n=64) and shifted to medium with 0.5 M sorbitol for 30-60 min (orange; n=62). Mean FRET ratio profiles together with 95% confidence intervals are shown.
